# A Spinal Circuit for Hypoxia-Evoked Motor Output in Zebrafish Larvae

**DOI:** 10.1101/2025.05.07.652688

**Authors:** Stephan Foianini, Kristian J. Herrera, Florian Engert

## Abstract

Oxygen availability is a critical environmental variable that shapes animal physiology and behavior. In larval zebrafish, acute hypoxia elicits a distinct increase in rhythmic pectoral fin movements, a behavior thought to facilitate oxygen uptake. While peripheral oxygen sensors such as neuroepithelial cells (NECs) and Merkel-like cells (MLCs) have been well characterized, the motor circuits responsible for executing this behavior remain unknown. Four distinct lower motor nerve branches in the spinal cord have been shown to innervate the pectoral fin muscles and are candidate effectors of hypoxia-induced behaviors. Here, we identify the neural pathways that transform hypoxia detection into a dedicated motor output. Using high-speed behavioral tracking, we confirm that hypoxia reliably increases pectoral fin beat frequency without affecting locomotor tail activity or visually guided swimming. Two-photon calcium imaging in Tg(ChaTa:Gal4;UAS:GCaMP6s) larvae reveals that a subset of cholinergic spinal motor neurons is selectively active during hypoxia-induced fin movements. Targeted laser ablation of pectoral fin motor nerves abolishes the response, demonstrating the necessity of descending input for this behavior. Our findings define a distributed, partially redundant motor circuit that implements a homeostatic fin response to hypoxia. By establishing a mechanistic framework for this behavior in a genetically accessible vertebrate model, this work enables future studies of oxygen sensing, sensorimotor integration, and the neural basis of homeostatic motor control.

## Introduction

Oxygen availability plays a crucial role in vertebrate physiology, influencing metabolic, cardiovascular, and behavioral responses^1–6^. In aquatic environments, hypoxia is a common and recurring challenge^7–18^, and many fish have evolved specialized adaptations to detect and respond to decreases in oxygen availability^19,20^. Among these adaptations, a distinct behavioral response observed in several species is the recruitment of pectoral fin movements during hypoxia^21,22^. These rhythmic, non-locomotor fin beats are thought to enhance oxygen uptake, possibly by disturbing the boundary layer around the gills or promoting fluid movement across respiratory surfaces. However, the prevalence of this behavior in larval fish remains incompletely understood.

Zebrafish (Danio rerio) larvae provide an ideal model to examine hypoxia-induced behaviors and their underlying neural mechanisms. Their optical transparency, genetic accessibility, and stereotyped motor behaviors offer a powerful platform for high-resolution functional imaging and manipulation^23–27^. Larval zebrafish also enable direct interrogation of the neural circuits that transform environmental stimuli into behavior, allowing for mechanistic insight into sensory-to-motor transformations that are difficult to access in other vertebrates. Although previous work has described fin movement responses to hypoxia in larval fish, the circuits driving this behavior have not been identified. As such, the larval zebrafish provides a rare opportunity to link a homeostatic sensory challenge to the specific motor neurons that implement a behavioral response.

At the sensory level, neuroepithelial cells (NECs) in the gills have been established as primary oxygen sensors in zebrafish, responding to hypoxia with serotonin release and synaptic communication to cranial sensory ganglia^28–30^. Merkel-like cells (MLCs), located in the oropharyngeal epithelium, have also been proposed as candidate oxygen sensors, although their functional roles are less well understood^31^. Previous work has shown that NECs can drive reflexive ventilatory behaviors, including changes in fin and jaw movements^32^, but how these sensory signals are transformed into coordinated motor output remains unknown.

In this study, we build on prior observations of hypoxia-induced pectoral fin movements in larval zebrafish by identifying the motor circuitry responsible for generating this behavior. Using high-speed behavioral tracking, two-photon calcium imaging, and targeted neural manipulations, we dissect the sensory-to-motor transformations that drive this response. We focus on characterizing the downstream motor components within the hindbrain and spinal cord that implement this oxygen-evoked behavior. By defining the circuit elements responsible for this response, we establish a framework for probing broader questions in oxygen sensing, sensorimotor integration, and the neural basis of homeostatic behaviors.

## Results

### Behavioral and Physiological Responses to Hypoxia

To determine whether larval zebrafish perform pectoral fin movements in response to acute hypoxia, we developed a behavioral assay to simultaneously monitor pectoral fin beats, tail movements, and heart rate in head-embedded 6 to 8 days post-fertilization (dpf) fish (Figure 1A). Localized delivery of hypoxic solution to the gills of the larva reliably induced rhythmic, non-locomotor pectoral fin movements that were distinct from baseline motor activity (Figure 1B).

**Figure 1:**
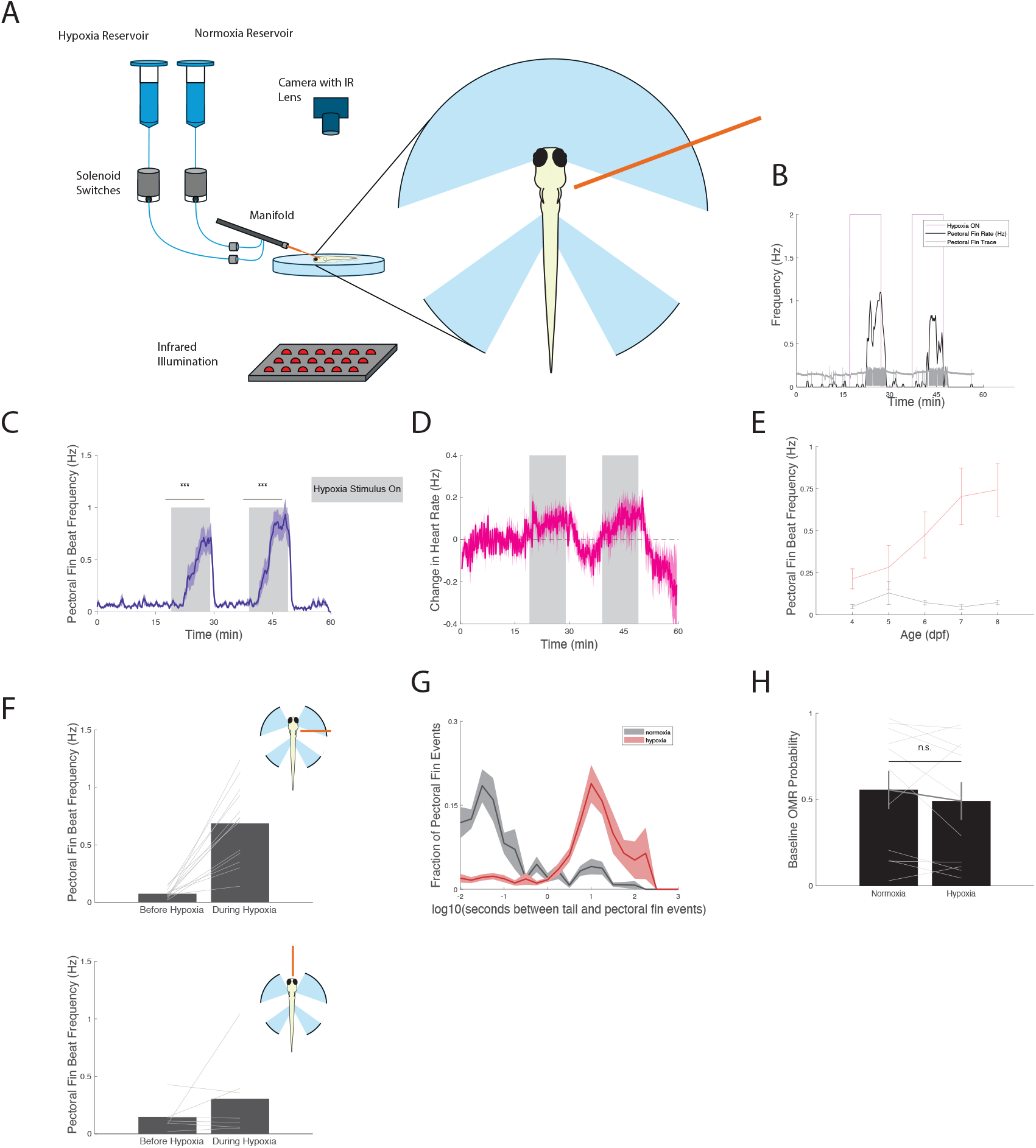
Effects of acute hypoxia exposure on larval zebrafish. A) Schematic of behavioral rig used to perform acute hypoxia exposure assays. B) Sample experiment showing pectoral fin behavioral readout and beat frequency of a single 7dpf fish. C) Pectoral fin beat frequency (Hz) of 6-8 dpf fish. D) Change in heart rate (Hz) normalized to a baseline taken before hypoxia stimulation. E) Change in pectoral fin beat frequency when fish are exposed to hypoxia throughout development. F) Pectoral fin beat frequency (Hz) during side and front delivery of hypoxia. G) Temporal relationship between tail and pectoral fin beats during normoxia and hypoxia. H) Probability of zebrafish performing an optomotor response (OMR) during normoxia and hypoxia.

Quantification across multiple fish revealed a significant increase in pectoral fin beat frequency during acute hypoxia compared to normoxia (Figure 1C). Notably, while heart rate increased almost immediately following the onset of hypoxia (Figure 1D), pectoral fin movements did not emerge until several minutes later, typically around the five-minute mark. This temporal offset suggests that distinct sensory thresholds or circuit mechanisms regulate cardiovascular and motor responses to hypoxia.

We next examined how this behavior changes across development (Figure 1E). In fish raised in normoxia, pectoral fin beat frequency in response to acute hypoxia gradually increased between 4 and 7 dpf, after which it appeared to stabilize. This developmental trajectory suggests that the fin motor program becomes progressively more robust as the underlying sensory and motor circuits mature. To test the spatial specificity of the hypoxia response, we delivered hypoxic solution either from the side (targeting the gills) or from the front (targeting the mouth). Fin movements were reliably evoked only with side-directed delivery, indicating that the gills are the relevant sensory site for this behavior (Figure 1F).

We next asked whether the increase in pectoral fin movements during hypoxia could be explained simply as a byproduct of swimming bouts, where pectoral fins are typically recruited alongside the tail during coordinated forward locomotion^33^. To address this, we examined the temporal relationship between pectoral fin and tail movement events by calculating the log-transformed time interval between each fin beat and the nearest tail beat (Figure 1G). During normoxia, the distribution was centered near zero, indicating that most fin movements occurred within a short window before or after a tail movement. This is consistent with pectoral fin activity being tightly coupled to bouts of swimming under baseline conditions. In contrast, during hypoxia, the distribution shifted significantly, with a much broader range of time delays between fin and tail events. The peak shifted away from zero, and a larger fraction of fin movements occurred independently of nearby tail activity. This decoupling suggests that hypoxia-induced pectoral fin movements represent a distinct motor program, rather than an extension of normal swim bouts. To ensure that the hypoxia-induced increase in pectoral fin movements did not result from a general disruption of motor function or sensory processing, we tested whether fish could still perform visually guided swimming during hypoxia. We used the optomotor response (OMR), a robust behavior in which larval zebrafish swim in the direction of moving visual stimuli to stabilize their position in the environment^26,34^. In our assay, we presented a pattern of dark, horizontally oriented stripes projected below the fish, which moved from tail to head, mimicking backward motion of the visual field. This reliably triggers forward swims as the fish attempts to compensate for perceived drift. We measured the probability of OMR-evoked swimming during normoxia and hypoxia and found no significant difference in response rates (Figure 1H). These results indicate that the sensory and motor systems required for vision-guided swimming remain intact during hypoxia exposure. Thus, the increase in pectoral fin movements during hypoxia is unlikely to reflect a nonspecific increase in motor drive or impairment in executing normal motor behaviors.

Together, these results demonstrate that larval zebrafish exhibit a specific pectoral fin motor response to acute hypoxia that emerges with a delay relative to cardiovascular changes. This behavior depends on hypoxia delivery to the gills and occurs independently of tail-driven locomotion.

### Neural Correlates of Hypoxia-Induced Behavior

To identify motor neurons involved in hypoxia-evoked pectoral fin movements, we performed two-photon calcium imaging in Tg(ChaTa:Gal4;UAS:GCaMP6s) zebrafish larvae, which express GCaMP6s in cholinergic neurons including motor pools. We focused our imaging on spinal cord segments and the region encompassing the motor root of the vagus nerve (cranial nerve X), which innervates the pectoral fins and heart, respectively^30,35^ (Figure 2A, top). Neural activity was recorded simultaneously with behavioral traces of fin movements, tail flicks, and heartbeats

**Figure 2:**
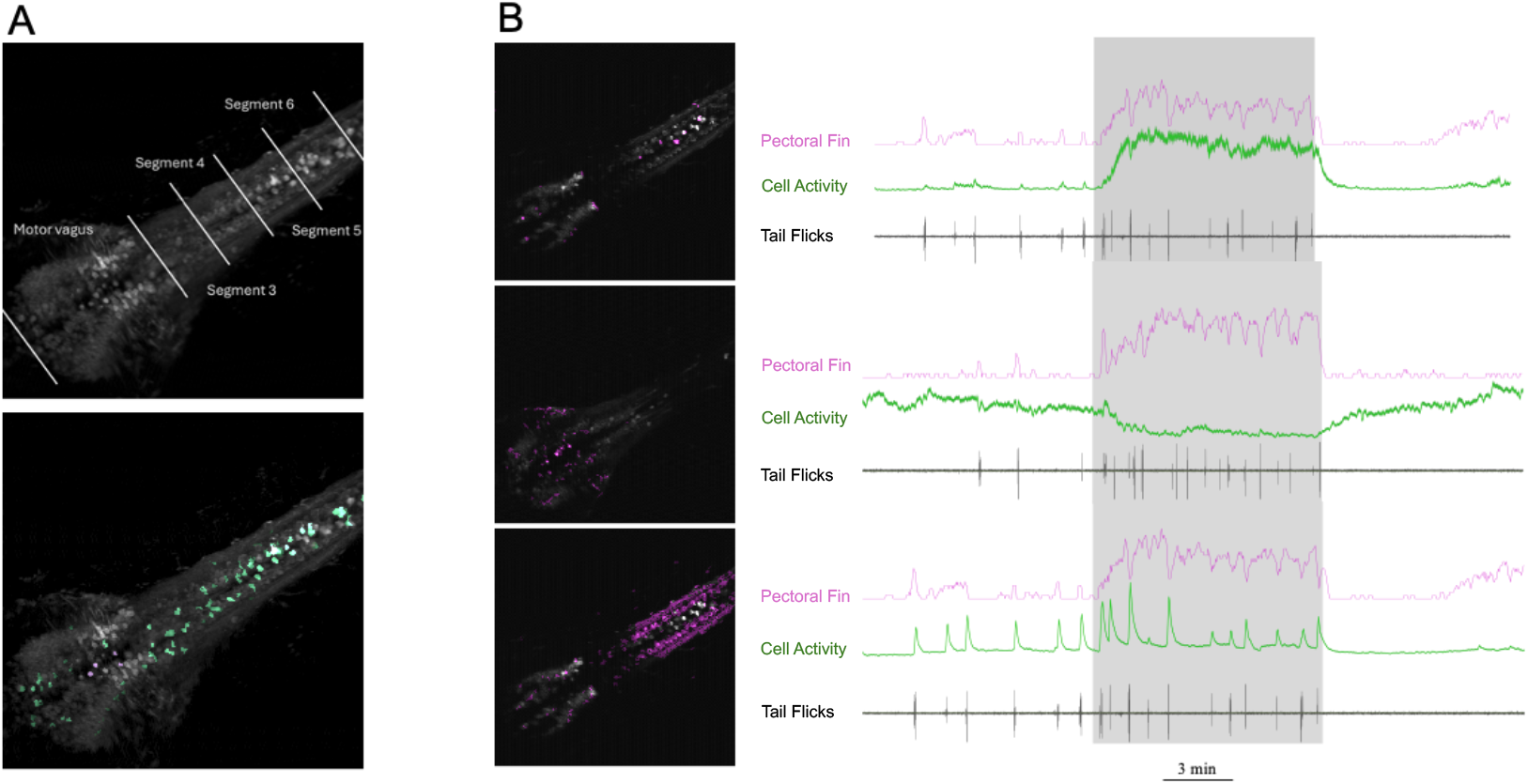
Calcium imaging motor neuron activity during hypoxia-induced pectoral fin movements. A) (Top) Image of a 7 dpf zebrafish highlighting the motor vagus nerve (cranial nerve X) along with the corresponding spinal cord segments that were imaged. (Bottom) Same image highlighting cells that show significant correlation with pectoral fin activity. Cells positively correlated with pectoral fin movements (**r > 0.8**) are marked in green, while cells that are anti-correlated with pectoral fin activity are marked in magenta. B) The left panels show cells active during pectoral fin movements (top), heart rate changes (middle), and tail flicks (bottom). On the right, the corresponding time traces illustrate the dynamics of pectoral fin activity (magenta), neuronal calcium activity (green), and tail flicks (black).

This analysis revealed distinct populations of neurons with activity patterns either tightly aligned or anti-correlated with fin movements. Cells with strong positive correlation to pectoral fin bouts (r > 0.8) were located primarily within specific spinal segments and are likely to represent fin motor neurons. In contrast, a separate group of neurons showed strong anti-correlation with fin activity (r < -0.8), suggesting reciprocal modulation within the motor network (Figure 2A, bottom).

To determine the specificity of these neuronal responses, we aligned activity from individual neurons to behavioral traces. Example cells showed clear increases in calcium activity that tracked closely with the onset and duration of pectoral fin movements, but not with heart rate changes or tail flicks (Figure 2B, top). Other neurons were selectively active during heart rate increases (Figure 2B, middle) or tail movements (Figure 2B, bottom), demonstrating that the responses of fin-related motor neurons are behaviorally specific. These results indicate that a subset of spinal and brainstem neurons are selectively recruited during hypoxia-induced fin movements, while others are either suppressed or unaffected.

### Redundant Motor Innervation Underlies Hypoxia-Induced Pectoral Fin Movements

To test the necessity of motor innervation for hypoxia-induced pectoral fin movements, we performed targeted laser ablation of pectoral fin motor axons in 6 to 7 dpf zebrafish. Pectoral fin muscles are innervated by four distinct cranial motor nerve bundles, which converge onto the fin base and are clearly visible under fluorescence microscopy^36^ (Figure 3A). We first confirmed that individual pectoral fin motor nerve bundles could be visualized and precisely targeted for ablation. In 6 dpf zebrafish, four distinct nerve bundles innervating the pectoral fins were clearly identifiable under fluorescence imaging. Using a two-photon laser, we selectively ablated individual bundles in vivo. Figure 3B shows a representative example: the left panel depicts a nerve bundle prior to ablation, with a red triangle marking the targeted ablation site. The right panel shows the same bundle immediately after laser ablation, with clear disruption of the axon tract, confirming the precision and effectiveness of the lesion.

**Figure 3:**
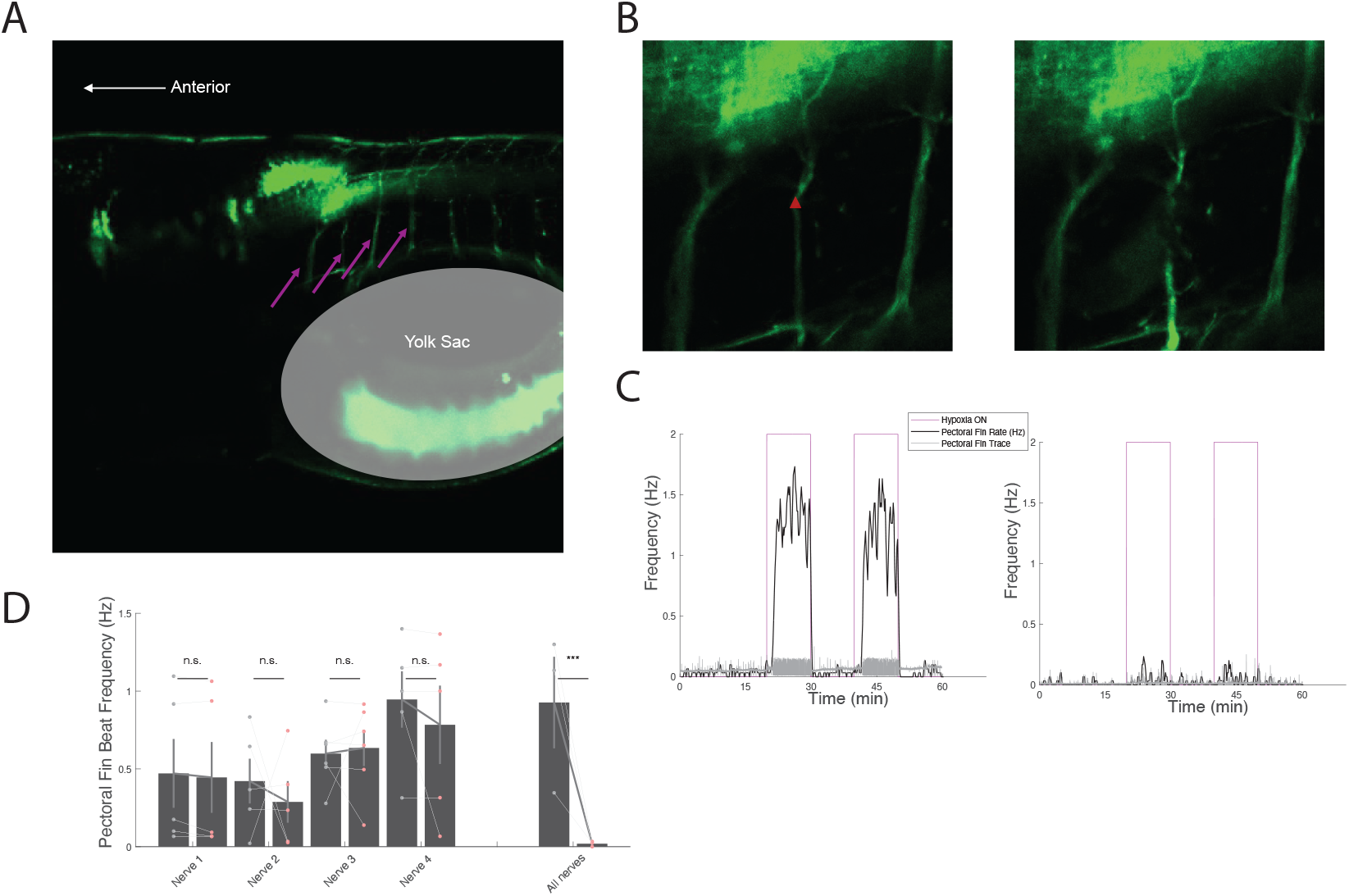
Laser ablation of pectoral fin nerve bundles reveals redundancy in neural control of hypoxia-induced fin movement. A) Lateral view of 6 dpf zebrafish pectoral fin innervation showing four distinct nerve bundles labeled with magenta arrows. B) Lateral view of a 6 dpf zebrafish showing an intact nerve bundle (left image) innervating the pectoral fin before laser ablation. Red triangle indicates the area ablated by the laser. Image on the right shows the same nerve bundle after laser ablation. C) Effect of complete unilateral motor neuron ablation on pectoral fin beat frequency during normoxia and hypoxia in a single 7dpf zebrafish. D) Effect of individual and complete nerve bundle ablation on pectoral fin movements during hypoxia in 7dpf zebrafish.

We then assessed the impact of ablating motor innervation on pectoral fin movements during hypoxia. In fish where all four motor nerve bundles on one side of the body were ablated, hypoxia failed to elicit fin movements on the lesioned side, while the intact side retained normal fin activity (Figure 3C). This result confirms that descending motor input is required to drive pectoral fin movement in response to hypoxia.

Next, we performed selective ablations of individual nerve bundles to test for functional redundancy in the circuit. Ablation of a single bundle did not significantly impair fin movement during hypoxia. However, when all four bundles were ablated, fin beat frequency was completely abolished on the lesioned side (Figure 3D). These findings indicate that the neural control of hypoxia-induced pectoral fin movements is distributed across multiple motor pathways, such that each bundle contributes partially and collectively ensures robust behavioral output.

## Conclusion

In this study, we define the motor circuit underlying a robust hypoxia-induced behavior in larval zebrafish. We show that pectoral fin movements are selectively and reliably recruited during acute hypoxia, independent of general locomotor activity. Through targeted calcium imaging in Tg(ChaTa:Gal4;UAS:GCaMP6s) fish, we identify cholinergic spinal motor neurons whose activity correlates with fin movement and is distinct from neurons involved in other motor outputs. Laser ablation of fin-innervating nerve bundles eliminates this behavior, confirming the necessity of descending motor input. These findings reveal a distributed, partially redundant motor circuit that converts hypoxia detection into a specialized motor response.

Although peripheral oxygen sensors such as neuroepithelial cells and Merkel-like cells have been characterized in zebrafish, the downstream pathways responsible for generating behavioral responses remain poorly understood. Our results bridge this gap by linking oxygen detection to motor execution at the circuit level. By uncovering the architecture of this response in larvae, we provide a mechanistic foundation for exploring how sensory information is integrated and routed through the brain to produce adaptive motor output.

The tractability of this behavior in larval zebrafish opens the door to mapping upstream sensory components, identifying integrative brain regions, and dissecting the neuromodulatory control of oxygen-responsive circuits. More broadly, our work establishes a framework for investigating how homeostatic stressors such as hypoxia are encoded, processed, and transformed into context-specific behavioral strategies.

Together, our findings establish larval zebrafish as a valuable model for uncovering how oxygen availability is transformed into behaviorally relevant motor output. By defining a specific and reproducible hypoxia-induced behavior and identifying the motor neurons required for its execution, we provide a foundation for future studies into the sensory, integrative, and modulatory components of this circuit. This work highlights how homeostatic challenges such as hypoxia can recruit distinct neural pathways to drive adaptive behaviors and demonstrates the utility of combining high-resolution imaging, targeted perturbations, and quantitative behavioral analysis in a genetically accessible vertebrate system.

## Methods

### Zebrafish Husbandry

Fertilized eggs were collected in embryo water containing Methylene Blue and E3 solution (5 mM NaCl, 0.17 mM KCl, 0.33 mM CaCl_2_, 0.33 mM MgSO_4_) and maintained at 28.5°C. After 24 hours, embryos were transferred to filtered facility water (200 nm pore size), which was refreshed daily to maintain water quality. Zebrafish were reared under standard conditions on a 14-hour light / 10-hour dark cycle. Beginning at 4 days post-fertilization, larvae were fed live paramecia. Unless otherwise noted, experiments were conducted on fish between 6- and 7-days post-fertilization. All experimental procedures adhered to institutional IACUC protocols as approved by the Harvard University Faculty of Arts and Sciences Standing Committee on the Use of Animals in Research and Teaching.

### Experimental Setup

Zebrafish larvae (7 dpf) were embedded in 2% low-melting agarose with their tails, pectoral fins, and gills free, following established immobilization protocols used in zebrafish neurophysiology studies. This setup ensured that the fish remained stationary while allowing natural fin and tail movements.

To induce hypoxia, oxygen levels in the hypoxic water reservoir were reduced using nitrogen bubbling, continuously monitored in real-time with an oxygen sensor and probe (PreSens Precision Sensing Fibox 4 trace/Oxygen Dipping Probe DP-PSt8). Oxygen concentrations were lowered to predetermined levels (2 mg/L dissolved O_2_).

Fluids were delivered via a gravity-driven system, directing flow precisely to the gills using a narrow, 360 mm diameter perfusion pencil tip (AutoMate Scientific 04-360) at a controlled speed of approximately 1.5 ml/minute. An 8-channel manifold (AutoMate Scientific 04-08-zdv) facilitated rapid solution exchange between hypoxic and normoxic water. Solution outputs were regulated by solenoids (Cole Palmer EW-01540-01) controlled via an Arduino®, ensuring that only one solution was presented to the fish at any time. Maintaining a continuous stream minimized behavioral responses to sudden flow changes and enabled rapid clearance of hypoxic water at the end of each trial.

### Behavioral Analysis

Pectoral fin movements were recorded using high-speed videography at 1000 frames per second to capture detailed kinematic features. Fin bouts were quantified using custom MATLAB scripts that tracked fin displacement and bout duration. Fin motion frequency and amplitude were compared between normoxia and hypoxia conditions.

To assess general locomotor function, we employed an optomotor response (OMR) assay. Fish were presented with a whole-field visual stimulus consisting of dark stripes drifting tail-to-head, projected from below. This reliably evokes forward swimming in larval zebrafish. OMR performance was measured under both normoxic and hypoxic conditions to ensure that hypoxia-induced fin movements were not due to global increases in motor drive.

### Calcium Imaging

To functionally identify neurons that were related to pectoral fin activity, we chose to perform calcium imaging using two-photon microscopy. Accordingly, we combined our gravity flow-based stimulus delivery system with a custom-built two-photon microscope as previously described^37^. During imaging, we recorded calcium activity plane-by-plane from a transgenic zebrafish that expressed GCaMP6s in all cholinergic neurons (tg(ChATa:gal4;uas:GCaMP6S^38^)). Imaging was targeted to the hindbrain and spinal cord, with every plane separated by 10 microns. Each plane was imaged for 30 minutes, where for the first 10 minutes of each plane the gills of the fish were exposed to normoxic water, followed by 10 minutes of hypoxic water, followed again by normoxic water. After imaging, calcium signals were extracted using previously described correlation-based segmentation^27,37^. These signals were then classified as being related either to tail or pectoral motion (or neither) based upon their correlation to regressors built from the convolution of either tail motion events or pectoral fin rates (Hz) with a GCaMP6s kernel.

### Neural Circuit Manipulation

Laser ablation experiments were used to test the necessity of descending motor innervation for pectoral fin movements during hypoxia. Fin-innervating motor nerve bundles were visualized under fluorescence microscopy and targeted using a Zeiss LSM 980 NLO multiphoton microscope equipped with a pulsed laser tuned for ablation. Individual bundles or all four bundles on one side of the body were selectively ablated. Successful ablation was verified by the visible disruption of nerve tracts and post-ablation behavioral responses were recorded under hypoxia to assess the loss of fin movement.

## References

1. Komniski, M. S., Yakushev, S., Bogdanov, N., Gassmann, M. & Bogdanova, A. Interventricular heterogeneity in rat heart responses to hypoxia: the tuning of glucose metabolism, ion gradients, and function. Am. J. Physiol. Heart Circ. Physiol. 300, H1645–H1652 (2011).

2. Chang, A. J., Ortega, F. E., Riegler, J., Madison, D. V. & Krasnow, M. A. Oxygen regulation of breathing through an olfactory receptor activated by lactate. Nat. Lond. 527, 240–244 (2015).

3. Thomson, A. J., Drummond, G. B., Waring, W. S., Webb, D. J. & Maxwell, S. R. J. Effects of short-term isocapnic hyperoxia and hypoxia on cardiovascular function. J. Appl. Physiol. 101, 809–816 (2006).

4. Richalet, J.-P., Hermand, E. & Lhuissier, F. J. Cardiovascular physiology and pathophysiology at high altitude. Nat. Rev. Cardiol. 21, 75–88 (2024).

5. López-Barneo, J. et al. Oxygen sensing by the carotid body: mechanisms and role in adaptation to hypoxia. Am. J. Physiol.-Cell Physiol. 310, C629–C642 (2016).

6. Siebenmann, C. & Lundby, C. Regulation of cardiac output in hypoxia. Scand. J. Med. 159.

7. Jenny, J.-P. et al. Global spread of hypoxia in freshwater ecosystems during the last three centuries is caused by rising local human pressure. Glob. Change Biol. 22, 1481–1489 (2016).

8. Dubuc, A., Waltham, N., Malerba, M. & Sheaves, M. Extreme dissolved oxygen variability in urbanised tropical wetlands: The need for detailed monitoring to protect nursery ground values. Estuar. Coast. Shelf Sci. 198, 163–171 (2017).

9. Blaszczak, J. R. et al. Extent, patterns, and drivers of hypoxia in the world’s streams and rivers. Limnol. Oceanogr. Lett. 8, 453–463 (2023).

10. Balangoda, A. Effect of Diurnal Variation of Dissolved Oxygen in a Eutrophic Polymictic Reservoir. Am. J. Environ. Sci. 13, 30–46 (2017).

11. Graham, J. B. Ecological, Evolutionary, and Physical Factors Influencing Aquatic Animal Respiration. Am. Zool. 30, 137–146 (1990).

12. Chang, W. Y. B. & Ouyang, H. Dynamics of dissolved oxygen and vertical circulation in fish ponds. Aquaculture 74, 263–276 (1988).

13. Jane, S. F. et al. Widespread deoxygenation of temperate lakes. Nature 594, 66–70 (2021).

14. Diaz, R. J. & Rosenberg, R. Spreading Dead Zones and Consequences for Marine Ecosystems. Science 321, 926–929 (2008).

15. Chapman, L. & Liem, K. PAPYRUS SWAMPS AND THE RESPIRATORY ECOLOGY OF BARBUS-NEUMAYERI. Environ. Biol. Fishes 44, 183–197 (1995).

16. Khan, A. A., Siddiqui, A. Q. & Nazir, M. Diurnal variations in a shallow tropical freshwater fish pond in Shahjahanpur, U.P. (India). Hydrobiologia 35, 297–304 (1970).

17. Jørgensen, B. B. & Revsbech, N. P. Diffusive boundary layers and the oxygen uptake of sediments and detritus. Limnol. Oceanogr. 30, 111–122 (1985).

18. Mandic, M. & Regan, M. D. Can variation among hypoxic environments explain why different fish species use different hypoxic survival strategies? J. Exp. Biol. 221, (2018).

19. Abdallah, S. J., Thomas, B. S. & Jonz, M. G. Aquatic surface respiration and swimming behaviour in adult and developing zebrafish exposed to hypoxia. J. Exp. Biol. jeb.116343 (2015) doi:10.1242/jeb.116343.

20. Kramer, D. L. & McClure, M. Aquatic surface respiration, a widespread adaptation to hypoxia in tropical freshwater fishes. Environ. Biol. Fishes 7, 47–55 (1982).

21. Jonz, M. G. & Nurse, C. A. Development of oxygen sensing in the gills of zebrafish. J. Exp. Biol. 208, 1537–1549 (2005).

22. Erickstad, M., Hale, L. A., Chalasani, S. H. & Groisman, A. A microfluidic system for studying the behavior of zebrafish larvae under acute hypoxia. Lab. Chip 15, 857–866 (2015).

23. Chen, A. B., Deb, D., Bahl, A. & Engert, F. Algorithms underlying flexible phototaxis in larval zebrafish. J. Exp. Biol. 224, jeb238386 (2021).

24. Howe, K. et al. The zebrafish reference genome sequence and its relationship to the human genome. Nature 496, 498–503 (2013).

25. Nasevicius, A. & Ekker, S. C. Effective targeted gene ‘knockdown’ in zebrafish. Nat. Genet. 26, 216–220 (2000).

26. Portugues, R. & Engert, F. Adaptive Locomotor Behavior in Larval Zebrafish. Front. Syst. Neurosci. 5, 72–72 (2011).

27. Portugues, R., Feierstein, C. E., Engert, F. & Orger, M. B. Whole-Brain Activity Maps Reveal Stereotyped, Distributed Networks for Visuomotor Behavior. Neuron 81, 1328–1343 (2014).

28. Pan, W., Scott, A. L., Nurse, C. A. & Jonz, M. G. Identification of oxygen-sensitive neuroepithelial cells through an endogenous reporter gene in larval and adult transgenic zebrafish. Cell Tissue Res. 384, 35–47 (2021).

29. Dunel-Erb, S., Bailly, Y. & Laurent, P. Neuroepithelial cells in fish gill primary lamellae. J. Appl. Physiol. 53, 1342–1353 (1982).

30. Jonz, M. G. & Nurse, C. A. Neuroepithelial cells and associated innervation of the zebrafish gill: A confocal immunofluorescence study. J. Comp. Neurol. 461, 1–17 (2003).

31. Pan, Y. K. & Perry, S. F. Developing zebrafish utilize taste-signaling pathways for oxygen chemoreception. Curr. Biol. 34, 4272–4284.e5 (2024).

32. Jonz, M. G., Fearon, I. M. & Nurse, C. A. Neuroepithelial oxygen chemoreceptors of the zebrafish gill. J. Physiol. 560, 737 (2004).

33. Green, M. H., Ho, R. K. & Hale, M. E. Movement and function of the pectoral fins of the larval zebrafish (Danio rerio) during slow swimming. J. Exp. Biol. 214, 3111–3123 (2011).

34. Naumann, E. A. et al. From Whole-Brain Data to Functional Circuit Models: The Zebrafish Optomotor Response. Cell 167, 947–960.e20 (2016).

35. Coccimiglio, M. L. & Jonz, M. G. Serotonergic neuroepithelial cells of the skin in developing zebrafish: morphology, innervation and oxygen-sensitive properties. J. Exp. Biol. jeb.074575 (2012) doi:10.1242/jeb.074575.

36. Thorsen, D. H. & Hale, M. E. Neural development of the zebrafish (Danio rerio) pectoral fin. J. Comp. Neurol. 504, 168–184 (2007).

37. Herrera, K. J., Panier, T., Guggiana-Nilo, D. & Engert, F. Larval Zebrafish Use Olfactory Detection of Sodium and Chloride to Avoid Salt Water. Curr. Biol. 31, 782–793.e3 (2021).

38. Förster, D. et al. Genetic targeting and anatomical registration of neuronal populations in the zebrafish brain with a new set of BAC transgenic tools. Sci. Rep. 7, 5230–11 (2017).

